# BADDADAN: Mechanistic Modelling of Time Series Gene Module Expression

**DOI:** 10.1101/2025.02.18.638670

**Authors:** Ben Noordijk, Marcel Reinders, Aalt D.J. van Dijk, Dick de Ridder

## Abstract

Plants respond to stresses like drought and heat through complex gene regulatory networks (GRNs). To improve resilience, understanding these is crucial, but large-scale GRNs (>100 genes) are difficult to model using ordinary differential equations (ODEs) due to the high number of parameters that have to be estimated. Here we solve this problem by introducing BADDADAN, which uses machine learning to identify gene modules—groups of co-expressed and/or co-regulated genes—and constructs an ODE model that predicts gene module dynamics under stress. By integrating time-series gene expression data with prior co-expression data it finds modules that are both coherent and interpretable. We demonstrate BADDADAN on heat and drought datasets of *A. thaliana*, modelling over 1,000 genes, recovering known mechanistic insights, and proposing new hypotheses. By combining machine learning with mechanistic modelling, BADDADAN deepens our understanding of stress-related GRNs in plants and potentially other organisms.

## Introduction

Climate change poses significant challenges to global food security due to its negative impact on crop growth and productivity (Suter and Widmer, 2013). With a growing world population, especially in climate-vulnerable areas (Dai, 2011), it becomes essential to develop crop varieties that are resilient to stresses such as pests, heat, and drought, particularly when these occur simultaneously or in rapid succession. To understand these plant environmental response mechanisms, as well as their potential trade-offs, we aim to understand the underlying gene regulatory networks (GRNs) (Quint et al., 2016), i.e. sets of genes which influence each other’s expression. The temporal dynamics of these GRNs are commonly modelled using ordinary differential equations (ODEs) (Cao et al., 2012; Valentim et al., 2015; de Luis Balaguer et al., 2017) fit to time series datasets. Importantly, such ODE models can reflect biological prior knowledge and enable causal insights through formal mathematical model analysis and human interpretation. They have successfully been used to describe the influence of a small group of genes on flowering time (Chew et al., 2022) or hypocotyl elongation (Nieto et al., 2022) in response to different temperature and light regimes, and enhanced our mechanistic understanding of these processes.

However, these models typically only include a small number of genes (<100) (Fröhlich et al., 2018) because they cannot (yet) be constructed for genome-scale plant stress response networks due to a lack of extensive, mechanistic, biological knowledge. On top of that, a large number of input genes makes parameter estimation prohibitively complex, as this requires significant amounts of data (Segal et al., 2003; Cao et al., 2012; Shaik and Ramakrishna, 2013) and results in exponentially more possible gene network topologies. To overcome these issues and enable ODE modelling of largescale regulatory networks from time series data, we focus on gene modules (also known as gene clusters or gene sets): groups of co-expressed and/or co-regulated genes (Segal et al., 2003; Saelens et al., 2018). By modelling temporal behaviour of gene modules instead of individual genes, we significantly reduce the number of interactions that require parameter estimates. Additionally, gene modules can help bridge the gap between single-gene expression levels and high-level phenotypes by linking these modules to broader biological processes or well-studied plant hormones. Finally, modules improve our ability to convert novel knowledge from model plants such as *A. thaliana* to agronomically relevant crops, since module-level insights are easier to translate between species than individual genes (Ficklin and Feltus, 2011; Julca et al., 2021; Shaik and Ramakrishna, 2013; Sreedasyam et al., 2023). Machine learning (ML) is a promising approach to extract gene modules from expression measurements, as it has demonstrated the ability to infer complex patterns from large amounts of (biological) data (Alber et al., 2019). Nonetheless, ML approaches do not elucidate interactions between biological mechanisms that underlie a plant’s response to environmental influences, and therefore fall short in delivering the detailed understanding that is needed to improve plant stress resilience. Thus, recently there has been an increasing interest in combining mechanistic models with ML approaches (Baker et al., 2018; Alber et al., 2019; Noordijk et al., 2024). In our context, this involves combining ML-derived modules, which extract relevant patterns from large amounts of biological data, with mechanistic models to provide interpretability and causal insights.

For insightful ODE modelling, we want to find modules that are both coherent and interpretable. Here, coherent means that genes within the same module share an expression pattern in a given dataset, allowing us to summarise their expression as a single variable (e.g., the mean) in our ODE model. Interpretable means that they can be linked to a clear biological process, providing insight into how plants respond to stress and deepening our mechanistic understanding. There is a trade-off between these two desired characteristics, which is related to the way in which co-expression is calculated in the context of gene clustering. Firstly, ‘local’ co-expression, calculated from gene expression measured within only one experiment (i.e. over various time points), might yield clusters that are coherent but overfit the data (Mao et al., 2019), resulting in an ill-defined biological function. On the other hand, ‘global’ co-expression is calculated from a gene expression compendium which covers a wide range of different conditions (e.g. developmental stages, tissues, or stresses). Global co-expression should reflect genegene relationships more generally, providing high biological functional similarity within modules, but potentially overlooking groups of genes that are only coexpressed in a particular context. In BADDADAN, we address this coherence-interpretability trade-off by employing a module finding approach that uses a combination of global and local co-expressions.

Module detection has been extensively researched and reviewed (Saelens et al., 2018), and has been used to infer functionally related groups of genes in various organisms from gene expression data. For example, modules were found by converting gene expression to an association network first, disregarding the temporal dimension (Langfelder and Horvath, 2008; Wang et al., 2008; Forster et al., 2022). Moreover, Janky et al. (2014); Mao et al. (2019) have extended such methods to incorporate prior biological knowledge. However, none of these methods explicitly leverage temporal data. In contrast, methods that exploit the temporal nature of the data (Heard et al., 2005; Schulz et al., 2012; McDowell et al., 2018; Han et al., 2021) do not build upon prior biological data, and suffer from sparse, noisy measurements, common in biology. Additionally, data integration has been employed to find modules (Heyndrickx and Vandepoele, 2012; Depuydt and Vandepoele, 2021; Forster et al., 2022; Sanchez-Munoz et al., 2024), but these methods cannot easily be applied to new transcriptomics datasets of interest, hampering their ability to improve gene module coherence. Finally, Lu et al. (2011); Wu et al. (2014); Soltanalizadeh et al. (2020) performed ODE modelling of gene modules in yeast, mice, or human, but did not build upon existing data, potentially reducing the interpretability of their modules and failing to leverage prior knowledge available for these species. Thus, many distinct elements of module finding, biological data integration, and ODE modelling are available, but no method yet integrates all these steps.

Here we present BADDADAN (a Bioinformatics Approach to Describe Dynamical Activations of a Dimension-reduced *A-priori*-informed Network) which integrates existing transcriptomics and transcription factor (TF) binding data with experiment-specific expression data to infer a dynamic network model of gene modules that are coherent and reflect clear biological functions. As a proof of concept, we show that we can quantitatively model gene module temporal expression through a system of ODEs with reasonable accuracy in two real-world datasets, and that these models recover known biological insight as well as provide novel hypotheses for drought or heat resilience mechanisms. Taken together, BADDADAN offers a quantitative and interpretable framework for studying temporal gene expression responses to external perturbations, applicable to plants and other organisms.

## Results

To demonstrate BADDADAN (Fig. 1), we apply it to two time series transcriptomics datasets in *A. thaliana* Col0: 1) a progressive drought stress dataset of 5-week-old plants grown in increasingly dry soil over a period of two weeks, with transcriptomics of leaf 7 sampled each day (Bechtold et al., 2016); and 2) a dataset where 6-weekold plants were subjected to elevated ambient temperature, with transcriptomic samples of rosette leaves collected unevenly over a 24 hour period (Caldana et al., 2011). We chose these datasets because they contain transcriptomics measurements over a large number of time points (>14), making them suitable for ODE modelling. Furthermore, they represent distinct stress response pathways, which allows us to test our approach on multiple types of stresses. Below, we will go over each step in our approach and its results on both datasets in more detail.

**Figure 1.**
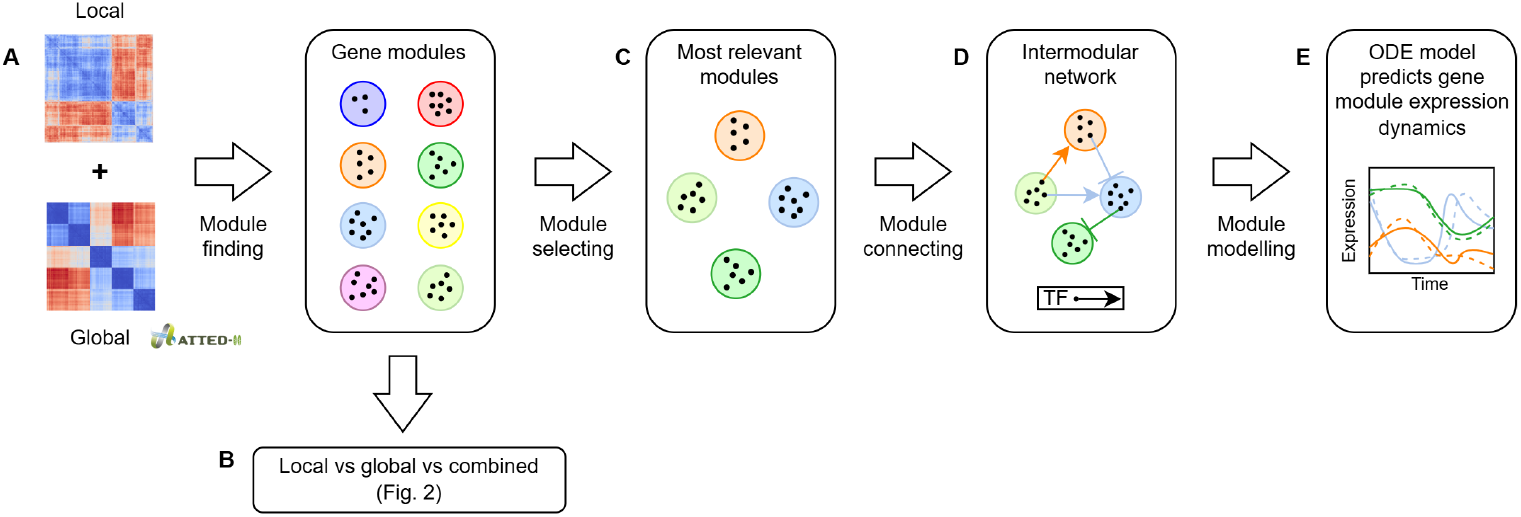
BADDADAN overview. (A)BADDADAN starts with candidate gene module finding based on a combination of experiment-specific (local) expression and compendium-wide (global) coexpression data (Obayashi et al., 2022). **(B)** To show this leads to improved coherence and interpretability, we compare these modules to modules created from either local or global data alone. **(C)** Our pipeline then selects a subset of the modules based on four criteria reflecting suitability for ordinary differential equation (ODE) modelling. **(D)** Next, it connects the modules using transcription factor binding site (TFBS) enrichment, and **(E)** creates an ODE model from this intermodular network which is fit to experimental (local) data and allows for biological insights.

### Module finding

As a first step of gene module finding, we filter our dataset for differentially expressed (DE) genes to reduce the chance of fitting modules to noise. DE filtering is performed using *limma* (Ritchie et al., 2015) (see Methods), and we find 2,363 and 4,444 DE genes (FDR*≤* 0.05) for drought and heat, respectively, with which we proceed for clustering. The set of DE genes is subsequently clustered. Hereto, we use either the (local) co-expression between the DE genes in the time-series data or (global) co-expression across a large compendium of datasets as compiled in ATTED (Obayashi et al., 2022), and also combine the local and global distances by summing them to express the similarities between the DE genes (see Methods). To investigate if clusters derived from the combined distance can balance coherence and interpretability, we compare them to clusters derived from either global or local distances alone. To assess coherence, we calculate the percentage of variance in gene expression over time explained by the first principal component for each module. To measure interpretability, we identify the enriched biological process gene ontology (GO) terms for each module (see Methods). Coherence is based on expression, which is used explicitly in the clustering process; in contrast, GO annotations are never explicitly used during the clustering, and thus provides an independent assessment of clustering performance. To prevent confounding (e.g. modules of size 1 are 100% coherent), we also ensure that the number of modules and their sizes are approximately equal between different clusterings (see Supp. Tab. S1, Supp. Fig. S1, and Methods).

In both datasets we see a similar pattern (Fig. 2): local distance-based clusters are more coherent than global distance-based ones, though fewer are enriched for at least one GO term. We furthermore find that combined distance-based modules balance coherence and interpretability, supporting our hypothesis that combining them can address the trade-off between the two. However, most differences between the use of local, combined, or global distances for the drought data are not significant (e.g. combined vs. local GO enrichment, *p* = 0.07, Mann-Whitney U test). This could be due to the smaller number of modules in drought (*≈*25) compared to heat (*≈*45), which, in turn, reflects the fewer differentially expressed genes in the drought experiment. The total number of GO terms per module follows a trend similar to the fraction of modules with at least one annotated GO term (Supp. Fig. S2). We also assess GO term functional relatedness per module using mean pairwise semantic similarity (Wang et al., 2007) (Supp. Fig. S3, Supp. Methods), showing that clustering on combined distances increases interpretability without reducing functional relatedness.

**Figure 2.**
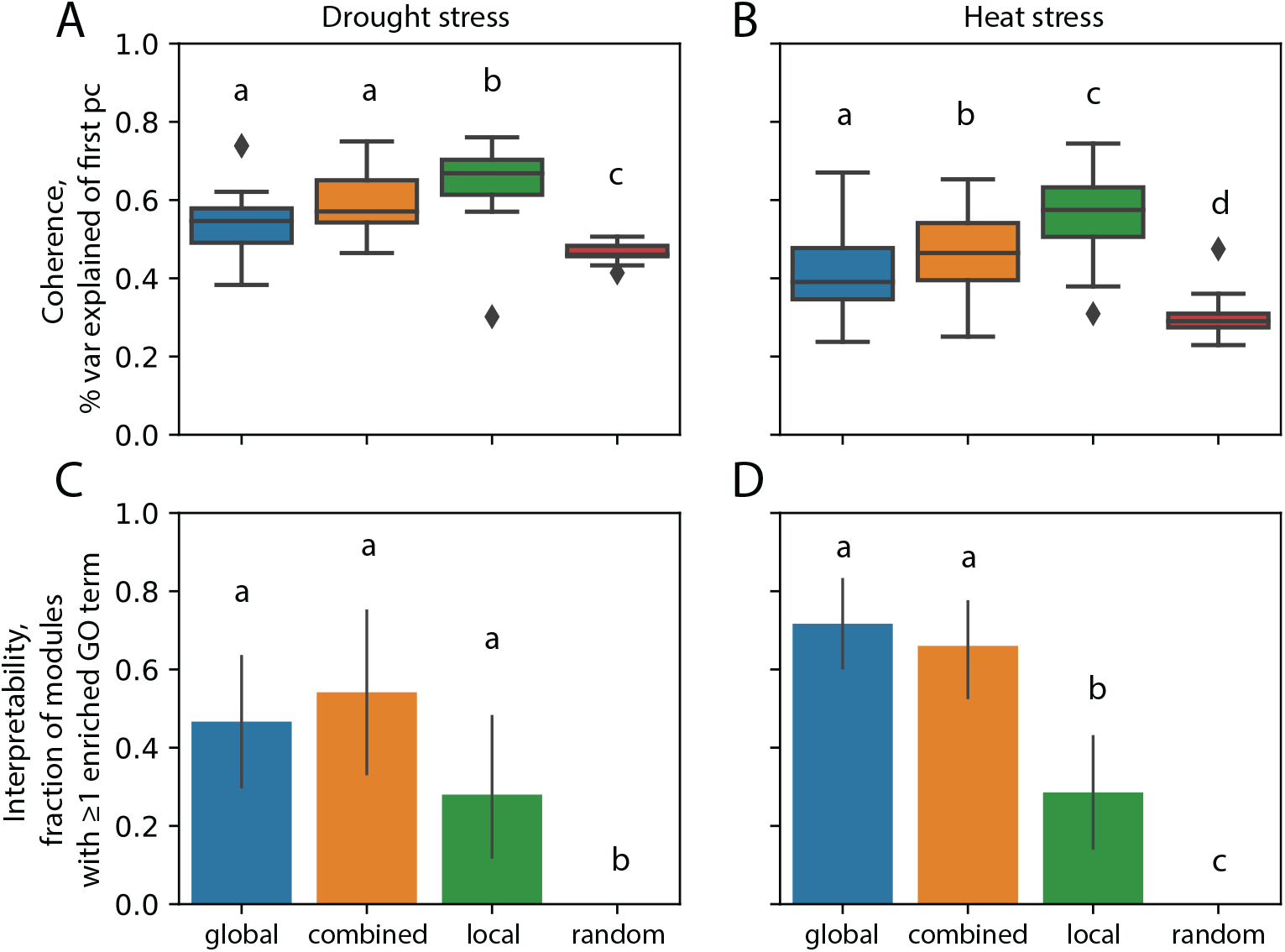
Module Finding. (**A,B**) Per-module coherence, i.e. explained variance of the first principal component of modules in drought and heat stress datasets, respectively. (**C,D**) Fraction of gene modules with at least one enriched biological process GO term. The x-axis represents the distances used to form modules: global (derived from compendium (Obayashi et al., 2022)), local (based solely on the experiment of interest), combined (a sum of local and global), and random (modules formed by random gene assignment). Error bars indicate the 95% c.i. estimated by bootstrapping. Different letters indicate *p <* 0.05, based on a two-sided pairwise Mann-Whitney U test between distributions.

### Module selecting

Not all found candidate modules may be relevant to include in our dynamic model. We select modules most suitable for ODE modelling based on four characteristics, expressed as z-scores (see Methods): 1) activity, i.e. a high average expression over all samples; 2) coherence, as this facilitates representation of the expression pattern by a single variable; 3) relevance, i.e. differing in expression between control and treatment conditions; and 4) regulatory relevance, hypothesising that modules with a high number of enriched transcription factor binding sites (TFBS) are more likely to be involved in stress response GRNs (Heyndrickx et al., 2014). There is no strong correlation between these four scores (Supp. Fig. S4, S5), suggesting their complementary nature. We use a combined Stouffer z-score of these four characteristics to proceed with the top 50% of all candidate modules (see Methods), resulting in 12 modules for drought and 26 modules for the heat stress datasets. A breakdown of the scores per module shows that the highest scoring modules score well on coherence, relevance, and activity, but not always on the regulatory relevance (Supp. Fig. S6).

### Module connecting

To incorporate a set of modules into an ODE model of intermodular regulation, we need to find regulatory interactions, i.e. determine which modules influence which other(s). Since such interactions cannot trivially be inferred from co-expression patterns and methods based on, for example, mutual information (Huynh-Thu and Geurts, 2018) do not always perform well (Aibar et al., 2017; Saelens et al., 2018), we choose to find these interactions based on existing knowledge of TFs and their TFBSs in *A. thaliana*. To this end, we used TF2Network (Kulkarni et al., 2018), which is based on extensive prior research (e.g. yeast one-hybrid and ChIP-Seq studies). However, connecting modules solely based on TFBSs can yield false positives, because TFs might not actually bind to a potential binding site due to e.g. low chromatin accessibility (Kulkarni et al., 2018; Khamis et al., 2018; De Clercq et al., 2021). Thus, we additionally require that a module can only be connected to another module when the TF that has a potential binding site to the target module is part of the source module and that this TF has an absolute correlation *>* .75 with the target module expression average (Supp. Fig. S7 and Methods). The sign of the correlation is then used to infer whether the TF activates or inhibits the target module. Moreover, we only consider a TF when the correlation between its expression and the average expression of its source module is above .3 (Supp. Fig. S8 and Methods). Finally, we only keep modules that have at least one ingoing or outgoing connection to other modules. The resulting intermodular networks are shown in Fig. 3b, and 3d for the drought and the heat stress network respectively.

**Figure 3.**
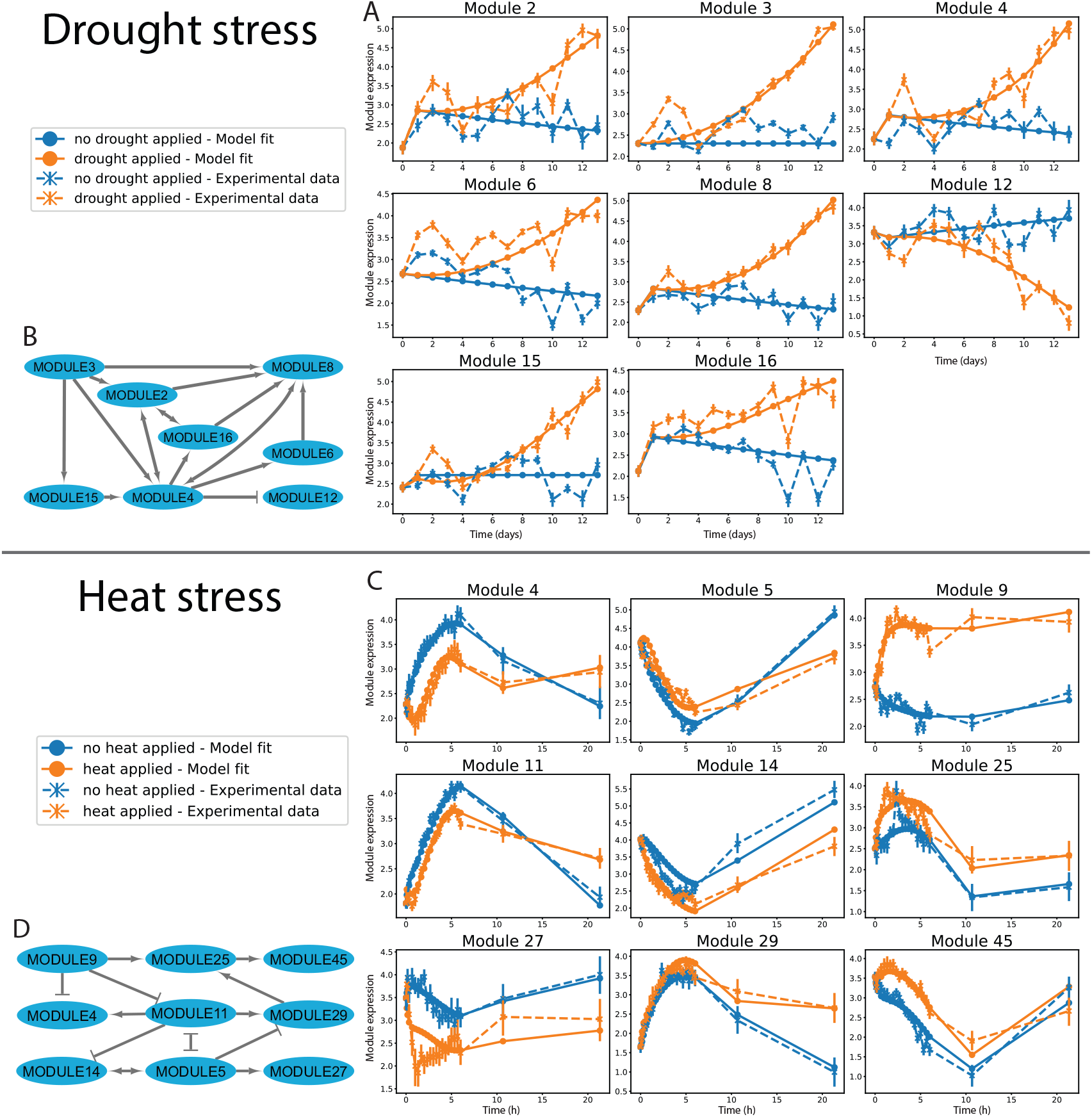
Intermodular networks and ODE fits. (**A,C**) Best ODE fit for the drought and heat stress experiments, respectively. Error bars indicate the 95% confidence interval of the mean module expression. (**B,D**) Intermodular network for drought and heat stress dataset, respectively. Arrows indicate activation, ‘T’-shaped ends indicate inhibition. Each edge can represent more than one TF. Module numbering is discontinuous due to the module selection step.

### Model fitting

Next, to evaluate whether BADDADAN is able to capture module expression dynamics, we construct ODEs based on the inferred module network, save them in SBML format (Hucka et al., 2003), and fit these using multi-start optimisation (see Methods). For the heat stress dataset, we explicitly model the circadian clock as input for the modules (see Methods) because the time points cover 24 hours and without the circadian clock as input our model was not able to attain an adequately accurate fit. For both datasets, the optimisation problem is hard; i.e. starting from 5,000 initial parameter sets, we rarely achieved the best objective value (Supp. Fig. S9).

Nevertheless, the resulting ODE models match the dynamics of the experimental data, with a mean squared error of 0.15 and 0.05 for drought stress (Fig. 3a) and heat stress (Fig 3c), respectively. However, they do not capture some of the early dynamics in the treatment condition (e.g. the peaks in module 3, 4, and 6 for drought stress or the dip of module 27 for heat stress). In the drought stress experiment, the conditions are expected to be highly similar between control and treatment for the first few days, so what causes these peaks is not *a priori* clear. The fact that early peaks are not fitted well could be due to the way the external input is modelled. For drought, this is a linear increase from 0 to 1 over 2 weeks. Consequently, the drought condition does not have a strong signal initially in the model. For the heat stress dataset, the external input into our model is a constant 1, even though the influence of the heat stress on certain modules might change over time (Kappel et al., 2023). Hence, further exploring the representation of the external input might enhance these fits. Alternatively, the fit may be improved by including additional module interactions or by adopting a more flexible ODE formulation (Eq. 7 in Methods).

### Mechanistic insights

We end up with ODEs for networks of eight and nine modules that represent 1,203 and 1,335 genes in total for drought and heat stress, respectively. Edges between modules are derived from one TF in most cases (12 edges for both drought and heat), with some edges representing two to six TFs (five and two edges for drought and heat, respectively). While comprehensive investigation of all interactions in these models is beyond the scope of this paper, below we select some hypotheses from our models and relate these to earlier work (Supp. Dataset S1, S2).

### Drought

For the drought stress experiment, the original publication (Bechtold et al., 2016) found that many genes only become differentially expressed from day eight onwards, as this likely marks a shift from mild to severe drought stress. We see a similar pattern in our data (Fig. 3a); many modules behave similarly in drought and control for approximately the first half of the experiment, and only diverge after that. Only two modules (25%) in our network could not be functionally annotated using GO terms (see Methods and Supp. Dataset S1), a far lower fraction than found in the original paper (Bechtold et al., 2016) where roughly 50% of the modules could not be annotated.

The interpretability of our approach helps generating hypotheses. For example, module 12, the only one with decreased expression levels in drought compared to control, is enriched for GO terms related to organismal-level ion homeostasis, iron ion starvation, and nutrient availability, suggesting that in the late phases of drought stress these can play a significant role, which has been found before (Mukarram et al., 2021; Mulet et al., 2020, 2023). According to BADDADAN, genes related to this homeostasis are inhibited by module 4 through the GBF3 (AT2G46270) TF, a bZIP G-box binding protein that has been found to be induced by water deprivation and was originally found in a finger millet drought transcriptome analysis (Ramegowda et al., 2017). Additionally, overexpression of this gene in *A. thaliana* has been shown to increase drought resistance (Ramegowda et al., 2017). As a potential mechanism, the authors suggest altered ABA signalling, but BADDADAN raises the possibility that GBF3 expression could be associated with ion homeostasis in drought resistance, which would be interesting to validate experimentally.

Module 4 and module 15 are enriched for terms related to oxidative stress and salicylic acid (SA) production or response, which are known to be involved in drought stress (Ortega et al., 2024). Analysis of the original paper’s data confirms that SA abundance correlates with the average expression of these modules in our ODE, with Pearson correlations of 0.61 and 0.66 for modules 4 and 15, respectively (Supp. Fig. S10). This demonstrates how modules can be successfully linked to higher-level phenotypic traits, such as plant hormone levels. SA has been shown to induce stomatal closure (Okuma et al., 2014), and there seems to be a negative correlation between stomatal conductance (Bechtold et al., 2016) and the expression of module 4 and 15. Unfortunately, given absence of the original raw measurement data, this is hard to quantify. Module 4 is also enriched for terms associated with biotic stress, in line with a role for stomatal closure in preventing water loss as well as bacterial entry (Okuma et al., 2014), and SA plays important roles in both biotic and abiotic stress (Ortega et al., 2024). Okuma et al. (2014) notes that stomatal closure is light dependent, which matches our model in that module 4 is regulated by module 2, which contains GO terms associated with light response. Module 4 has the highest closeness centrality of 1 in the network, which suggests it is more likely to play a key role in the overall response mechanism. Finally, module 16 is regulated by both module 2 and 4 and enriched for starch, glucan, and maltose metabolism. This suggests that these metabolic processes could be controlled through, e.g., circadian rhythm and light—all from module 2—but also stress-related processes from module 4, all of which have been reported (Lu et al., 2005; MacNeill et al., 2017; Skryhan et al., 2018). Knocking out *in silico* (see Methods) the CBF3 (AT4G25480) TF that connects module 2 to module 4 results in altered expression of e.g. module 16 (Supp. Fig. S11), and in literature it is indeed reported that changed CBF3 expression can impact sugar metabolism (Gilmour et al., 2000). All in all, these findings show that BADDADAN recovers known mechanisms and TFs, and can link module expression to other measurements such as plant hormone abundance.

Note that the expression profiles of some modules appear relatively similar (e.g., those of module 2 and 4). Recall that the modules are created by clustering both the local distances as well as the global distances that came from a compendium of data. The latter reflects that the two modules will have different biological functions. This is indeed true as the GO enrichments of modules 2 and 4 are different; module 2 is related to light response, circadian rhythm, and organic hydroxy compound biosynthetic process, whereas module 4 is related to biotic and abiotic stress. This shows the potential benefit of clustering based on a combination of local and global expression measurements.

### Heat

The heat stress dataset shows a large variety in up- and downexpression of modules over time, which is reflected in a large number of edges representing inhibition in the intermodular network (six inhibiting edges and eight activating ones, vs. one inhibiting and 16 activating in drought).

Module 4 displays interesting dynamics, initially lower in heat than control but higher at the last time point. Module 4 is linked to heat and temperature stress responses as well as protein folding. Potentially, module 4 might be involved in longer-term thermotolerance, as found recently for one of its constituent genes (HSFA2) (Pan et al., 2024). BADDADAN suggests this is potentially through protein-folding-related processes, evidence for which can be found (Wang et al., 2024). The expression pattern of module 4 seems to result from the type-2 coherent feed-forward loop (Alon, 2007) from module 9 through module 11. Module 9 inhibits module 4 through the LCL1 (AT5G02840) TF, which has been found to be involved in heat shock response (Kidokoro et al., 2023). Module 11 stimulates module 4, 5, 14, and 29 through the ABF1 (AT1G49720) TF, which is linked to ABA response as well as heat (Song et al., 2017), highlighting it as a potential key regulator. Unfortunately, ABA was not measured in this experiment, so we cannot study how its abundance correlates to module expression. Nonetheless, Knocking out ABF1 *in silico* (see Methods) reveals a stronger influence on modules 4 and 29 compared to modules 5 and 14 (Supp. Fig. S11). Specifically, the parameters *β*_11,4_*≈* 195 and *β*_11,29_*≈ ·* 20 are much higher than *β*_11,5_*≈* 7*·*10^*−*4^ and *β*_11,14_*≈* 4*·*10^*−*4^, highlighting that not all inferred intermodular edges represent equally strong relationships. This underscores the value of the interpretable parameters provided by our approach.

Module 5 is initially downregulated in both conditions, with a slightly higher expression in heat in the first 10 hours, after which we observe a higher expression in the control condition. This module is associated with defence response, stomatal movement, and response to biotic (bacterial) stimulus. As we also found in the drought dataset, the biotic stimulus is likely related to the stomatal movement (Okuma et al., 2014). Module 5 is also enriched for programmed cell death, glucosinalate catabolism, and callose localisation, which have been linked to (a)biotic stress responses (Hopkins et al., 2009; Janda et al., 2019). Overall, this shows how BADDADAN leverages prior knowledge to identify modules potentially related to combined stresses, and suggests that module 5 is an interesting target for studies on interactions between them.

Finally, Module 29 exhibits a similar response in control and heat during the initial phase of the heat shock, but at later time points is more active in heat. This could again be through a type-2 coherent feed-forward loop (Alon, 2007), this time from module 5. Module 29 is related to cellular homeostasis, glucosinolate transport, and sulphur compound transport, all related to heat stress (Sugio et al., 2009; Ihsan et al., 2019). An increase in catabolism of glucosinalate (which contains sulphur) due to module 5 will lead to a need for transport. Module 5 inhibits module 29 through the LHY1 (AT1G01060) TF, linked to the circadian clock; BADDADAN suggests that it could also be involved in controlling sulphur transport, through module 29. This has been observed in *Medicago truncatula* (Achom et al., 2022), and LHY1 has been linked to sulphur in *A. thaliana* (Van De Mortel et al., 2008). As the underlying mechanism is still unknown, this is a potentially interesting direction for future research.

Of the modules that initially display a large difference between the control and heat conditions (4, 9, 25, 27, 45), only three could be annotated for function (4, 9, 25). Module 9 has interesting annotations such as fluid transport—which suggests the early phase response of heat is causing the plant to transport more water—and hydrogen peroxide transmembrane transport, but because it has no ingoing connections we cannot infer what processes or TFs are involved in regulating this. This shows that BADDADAN only recovers a part of the intermodular network and thus does not necessarily provide insight into the full network; however, any model based on a subset of genes will be similarly limited.

## Discussion

Here, we introduce BADDADAN, a framework for constructing an ODE model of gene modules encompassing more than 1,000 genes in total. We apply it to two real-world datasets covering different stresses and time scales, demonstrating its advancements in gene module expression modelling. First, by combining TFBS enrichment with expression data, it identifies directional, signed relationships between modules. Second, by estimating parameters from data that capture these relationships, it quantifies interaction strengths. Third, by combining experiment-specific expression data with compendium-wide co-expression data, it yields coherent and interpretable modules. Together, these features enable accurate predictions of gene module expression over time and *in silico* predictions of responses to perturbations, such as knockouts, that would not be apparent from network structure alone. Moreover, BADDADAN uncovers known biological processes and their interactions while also generating novel hypotheses that can inform the design of future experiments.

BADDADAN enables out-of-the-box integration of experiment-specific data with prior experimental data (ATTED (Obayashi et al., 2022)), addressing a limitation of the widely-used WGCNA (Langfelder and Horvath, 2008), which does not directly support such integration and assumes a scale-free network topology. In contrast to recent black box machine learning approaches (Sapoval et al., 2022; Baker et al., 2018; Shen et al., 2021), BADDADAN is composed of intermediate steps that are inspectable; e.g., module selection provides z-scores, connections between modules can be visualised as a directed graph, and the ODE model contains human-interpretable parameters (e.g. *β* can be linked to interaction strength). Moreover, existing ODE modelling approaches comparable to BADDADAN (Lu et al., 2011; Wu et al., 2014) infer connections between modules solely in a data-driven way—similar to LASSO regression—which has been shown to underperform for GRN inference (Saelens et al., 2018). However, as potential future research, such connections could be integrated with our TFBS-based ones, a hybrid approach similar to Aibar et al. (2017) where connections are inferred if they are important for predicting dynamics and/or underpinned by the presence of a TFBS.

By addressing essential challenges in gene module network modelling—such as edge directionality determination, activation/inhibition inference, and associated weights estimation within an ODE framework— BADDADAN opens up extensive possibilities for further analysis. For example, looking deeper into confidence interval estimation of parameters, applying an identifiability analysis to our model, or performing topological sensitivity analysis (Babtie et al., 2014), where multiple different network topologies are tested for their ability to describe the data. This will tell us if we have found a network topology which would uniquely describe the resulting behaviour. Related to this, approaches for optimization of a given network topology by adding or removing a small number of nodes (modules in our case) could be applied (Astola et al., 2014).

Settings of each individual step are now optimised separately in the pipeline, but it would be an interesting to research if gains can be made by optimising the pipeline end-to-end through differentiable optimisation methods developed for deep learning (AlQuraishi and Sorger, 2021). This would also allow the use of deeplearning-based embeddings for clustering, higher dimensional representations of module expressions, and neuralODEs or biologically informed neural networks for ODE fitting. On top of that, the loss function could be extended to make modules adhere to e.g. rate limits or other existing biological knowledge.

Furthermore, to aid in integration with existing or novel methodologies, BADDADAN builds on existing standards and software. For parametrisation of the ODE we utilise standard formats and scripts for systems biology models (Schmiester et al., 2021; Schälte et al., 2023), and we save our our models as SBML format files (Hucka et al., 2003), allowing interaction with the wide ecosystem that is already available for systems biology models. This would allow us, for example, to couple BADDADAN to high quality ODE models for small GRNs that are already available (Chew et al., 2022; Valentim et al., 2015; Nieto et al., 2022). Given that both are comprised of ODEs, integration should be possible. For example, we could use a BADDADAN module related to the production of a metabolite—similar to how we linked two modules to SA in the drought dataset— and use the abundance of that metabolite predicted by BADDADAN as input for an existing mechanistic model. Vice versa, mechanistic models that predict metabolite abundance could be used as input (cf. *c*(*t*) or *u*(*t*)) to BADDADAN. Alternatively, a TF in the BADDADAN model could be taken to directly influence expression of a gene in the mechanistic model, improving its prediction accuracy.

Finally, in the future BADDADAN could be applied to datasets of other (non-)plant species and/or stresses and could form the basis for translational science, using fundamental insights into model organism *A. thaliana* resilience mechanisms to breed relevant crops with higher resilience. Since our model can predict possible KO effects under different stresses it also provides possibilities to study the effects of genetic variation on resilience. Also, a comparative analysis of modules between different individual or combined stresses could give insight into the temporal dynamics of processes important for multiple stresses specifically (Rizhsky et al., 2002; Verslues et al., 2023), which would further support targeted resilient crop development.

Summarising, BADDADAN provides a proof of concept approach for inferring gene module-based dynamic models from time series transcriptomics data, to help unravel the intricate gene regulatory mechanisms that plants employ to deal with stresses. Increasing knowledge in this area might eventually be used to breed more climate resilient crops which can be used to provide food to a growing world population, even under the pressure of climate change.

## Methods

### Gene expression input data

Progressive drought microarray expression data (Bechtold et al., 2016) was extracted from Gene Expression Omnibus (GEO) (Barrett et al., 2013) accession GSE65046, and probe names were converted to TAIR gene IDs using the Python package GEOParse (v2.0.3) (Gumienny, 2024). For the heat dataset (Caldana et al., 2011), expression data was extracted from ArrayExpress accession E-MTAB-375 and again probe names were mapped to gene identifiers using the annotations available on GEO (Barrett et al., 2013). Both original datasets were already log_2_-transformed to make noise additive instead of multiplicative and to stabilise variance (Archer et al., 2004).

### Differential gene expression

We used *limma* (v3.60.3) (Ritchie et al., 2015) in R (v4.4.0) for differential gene expression analysis, as it has been shown to perform well on time-course microarray datasets (Moradzadeh et al., 2019). For the drought stress dataset, contrasts were used to test if genes changed expression differently between subsequent time points in the drought group compared to the control group. However, such contrast testing only works if multiple biological replicates are available, which was not the case for the heat stress data. Thus, we ran *limma* on this dataset by fitting a natural cubic spline with five degrees of freedom through the time points using the R *splines* library. Next, we used *limma* to conduct an F-test between the conditions on the five parameters that correspond to interaction, i.e. difference between the control and heat group. Finally, for both drought and control, only genes with a BenjaminiHochberg false discovery rate (FDR) *≤*0.05 were kept for further processing.

### Module inference

To obtain global (compendium-wide) co-expression patterns, we downloaded pairwise z-scores from ATTED v11.1 (Obayashi et al., 2022), which were calculated based on 27,427 samples and corrected for biases due to differences in sampling conditions and sample sizes (Obayashi et al., 2022). The z-scores (denoted with *W*_global_) were converted to a distance matrix *D*_global_, which represents pairwise distances between genes, with each row and column corresponding to a gene:

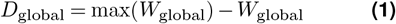

where max(*M*) = max_*i,j*_ *M*_*ij*_ is the maximum value over all rows and columns. This is consistent throughout the manuscript and also holds for uses of min. Local distances were calculated within each dataset as follows:

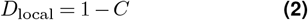

where *C* is the matrix of pairwise Pearson correlations between genes over the samples (taking the average expression level if biological replicates were available). Saelens et al. (2018) found that this ‘unsigned’ (Langfelder et al., 2011) metric yielded higher module detection accuracy compared to e.g. an absolute Pearson correlation or Spearman correlation.

To obtain a clustering based on both global and local coexpression, we combined *D*_global_ and *D*_local_. To avoid biases due to a difference in distributions, we first separately converted both distances to a z-score

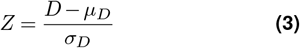

where *µ*_*D*_ is the average and *σ*_*D*_ the standard deviation of all distances.

Next, we summed the resulting z-score matrices:

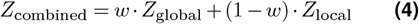

where *w* influences the balance between global and local co-expression. We set *w* = 0.5 for all experiments, but *w* could be set differently for specific purposes; e.g., assigning a higher weight to local co-expression if coherence is desired.

Finally, to ensure no distances were negative (needed for clustering), we converted them to a positive distance:

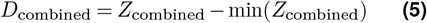

To find modules, we use hierarchical clustering. This has been shown to obtain reasonable performance in gene module detection (Saelens et al., 2018), where it was benchmarked on simulated and experimental data with a known ground-truth. Average linkage hierarchical clustering was performed in R on *D*_combined_, *D*_global_, and *D*_local_. To extract modules from the resulting dendrogram, it could be cut at a specific height, but such a global approach has been shown to be suboptimal for gene module detection (Langfelder et al., 2008). Thus, we applied a Dynamic Tree Cut method (Langfelder et al., 2008) designed specifically for gene module detection. Also, since module size affects coherence (e.g. modules of size 1 are 100% coherent), we ensured that module sizes were roughly equal between the three clusterings by changing the deepsplit parameter which influences cluster size (Supp. Fig. S1). For the drought stress data, we set deepsplit=1 for the global and combined distance matrices, but used deepsplit=2 for the local one, as this ensured cluster sizes similar to the other two clusterings (Supp. Fig. S1a). For the heat dataset, module sizes were comparable for all input distances at deepsplit=1 (default value) (Supp. Fig. S1b). All other Dynamic Tree Cut settings were kept at their default values.

### Characteristics of clusterings

We compared our local, combined, and global distancebased clusterings on coherence and interpretability (GO enrichment) of the resulting modules.

### Coherence

To assess coherence, we performed a PCA on each module separately, where we first normalised the expression of each gene to have mean 0 and variance 1 (Eq. 3). Coherence was then calculated as the percentage of variance explained by the first principal component, found using the PCA function in Scikit-learn (v1.1.3) (Pedregosa et al., 2011). We plotted the distribution of these coherence scores over all modules using Seaborn (v0.11) (Waskom, 2021).

### GO enrichment

GO enrichment was performed using GOAtools (v1.2.3) (Klopfenstein et al., 2018) with all differentially expressed genes as background. Only gene annotations of experimental evidence codes were used to prevent data leakage from existing co-expression-based electronic annotations. We focused on biological process GO terms, as we were interested in biological interpretation of the gene modules. For all modules, only enriched GO terms with a Benjamini-Hochberg FDR*≤* 0.05 were used for further analysis. To study the variability in the fraction of modules that had at least one enriched GO term, we used Seaborn’s bootstrapping (Waskom, 2021) to estimate the 95% confidence interval.

### Statistical testing

Statistical tests were performed using the Python package statannotations v(0.5.0) (Charlier et al., 2022). Because we could not assume all data to be normally distributed, we compared coherence and interpretability using the non-parametric two-sided Mann-Whitney U test. As a null model we measured the characteristics of random modules, where we ensured that the module sizes were distributed identically to those of the combined distances by randomly permuting these cluster assignments.

### Module selection

To select candidate modules for modelling, we assigned each module a score composed of four different components. 1) Activity: to obtain modules that are composed of genes that are active, we determined the average expression over all samples of all genes in a module. 2) Coherence: to measure the coherence of a module, we performed PCA on the normalised expression of its constituent genes over time in our dataset of interest and measured the percentage of variance explained by the first principal component, as described above. Coherence is desired because it facilitates representation of the module expression by a single value (e.g. average). 3) Relevance: to assess whether a module’s expression differs between treatment and control, we first summarised the module by calculating the average expression of its genes over time for each condition. We then measured the difference between conditions using the mean squared error of these averaged expression profiles. 4) Regulatory relevance: we assumed modules with a high number of enriched transcription factor binding sites (TFBS) were more likely to be important for a plant’s stress response; e.g. genes related to stimulus response and gene regulation in *A. thaliana* are known to have a higher number of regulators (Heyndrickx et al., 2014). Moreover, modules with many enriched TFBS are more likely to successfully be linked to other modules in our model. To score this regulatory relevance, we used the count of TFBS *N*_*T*_ that were enriched in a module according to TF2Network (Kulkarni et al., 2018). Since this distribution was highly skewed, we applied a transformation log(*N*_*T*_ + 1). We chose not to include interpretability (i.e. GO enrichment) in the scoring to ensure that it remains an independent criterion for evaluating the selected modules.

All scores were converted to z-scores (through Eq. 3), and subsequently summed using Stouffer’s method to yield a final summary score for each module:

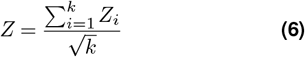

Here, *k* is the number of different characteristics (in our case, four). Only the activity was based on non-normalised gene expression, the other scores were calculated from gene expression where each gene was zscored over all samples. We selected the top 50% (a user parameter, here chosen arbitrarily) of all modules based on their summary score for subsequent modelling.

### Model structure

To infer connections between modules, we used the TFBS enrichment results of TF2Network (Kulkarni et al., 2018). We inferred regulatory interactions between modules based on three criteria: 1) The target module contained an enriched binding site for a TF that belonged to a source module. 2) The TF had a Pearson correlation of at least 0.3 with the average expression of its source module; any TFs with correlations to their source module lower than this were considered an outlier and excluded from regulatory interaction inference. 3) The absolute Pearson correlation between the expression of the TF and the average expression of its target module had to exceed a certain threshold. This step fulfilled two functions. Firstly, it reduced the number of false positives that could result from TFBS enrichment analysis, e.g. due to chromatin accessibility (Kulkarni et al., 2018; Khamis et al., 2018; De Clercq et al., 2021). Secondly, as we needed to choose a different form of the Hill equation for inhibition of activation (Eq. 7), we used the sign of the Pearson correlation as a heuristic to infer up- or downregulation. If multiple TFs from the same source module regulated the same target module, we asserted they all consistently upor downregulated; if there was disagreement, no valid network would be output by BADDADAN. Finally, modules that were left with no interactions with any other modules were removed from the model.

To determine the correlation cutoff between TFs and their target modules, we inferred model structures as described above for cutoffs varying from 0 to 0.95 at intervals of 0.05, and kept track of the total number of modules, interactions, and whether the model graph consisted of one single weakly connected component or multiple disjoint components (Supp. Fig. S7). We then picked 0.75 as a cutoff for both the heat and drought stress dataset, as this led to the sparsest model not split into disjoint components. Intermodular network visualisation and analysis were performed in NetworkX (v2.8.8) (Hagberg et al., 2008) and Cytoscape (v3.10.3) (Shannon et al., 2003).

### ODE model

For each module, we model its activity in an ODE representing the average gene expression levels over time after normalization (Eq. 3) because related genes can have similar expression patterns but at different magnitudes (Wu et al., 2014). Since z-score normalisation of the gene expression could result in negative values, a constant of 3 (minimum value of module expression was around −2.5) was added to guarantee positive values, preventing biologically implausible scenarios where negative expressions would invert the decay term and falsely suggest gene production. For a module *m* that is inhibited by a set of modules *I* and activated by a set of modules *A*, we represent its rate of change as follows:

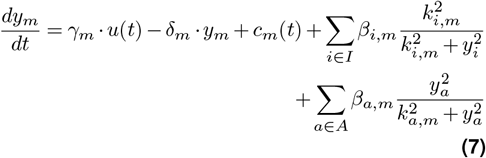

The expression of each module *y*_*m*_ is dictated by an external stimulus-dependent term weighted by *γ*_*m*_, a decay term governed by the parameter *δ*_*m*_, a circadian clock input *c*_*m*_(*t*) that we only used for the heat stress dataset (because this is sampled over a period of 24 hours), and activation/inhibition Hill equations of the second order where *β*_*n,m*_ represents the maximum magnitude of the activation/inhibition of module *n* on module *m*, and *k*_*n,m*_ the abundance of the regulator at which the module is 50% activated/inhibited. We chose these formulas as gene regulation is highly nonlinear and Hill kinetics are widely used in ODE modelling in systems biology (Polynikis et al., 2009; Frank, 2013; Valentim et al., 2015). The function *u*(*t*) captures the external input, which was different for the drought and heat experiment. For the former, we modelled the linear increase in drought stress over two weeks (Bechtold et al., 2016) as follows:

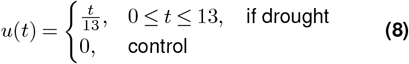

to simulate a linear increase of drought where *t* is time in days.

For the latter, heat stress, we modelled the external input as follows:

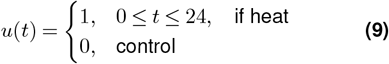

as is also common in plant mechanistic modelling papers (Nieto et al., 2022). Here, we measured *t* in hours. Finally, because initially the model could not adequately model the data in the heat experiment, we explicitly added a circadian clock *c*_*m*_(*t*) as input for each module:

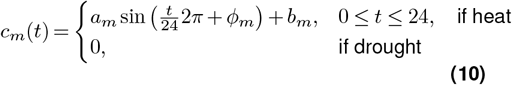

Here, *a*_*m*_ quantifies the input strength of the circadian clock on the module expression change, *ϕ*_*m*_ determines the phase offset of the circadian clock influence (which we give a 24h period by definition) and *b*_*m*_ models the baseline influence of the circadian clock. Again, *t* is measured in hours. Finally, BADDADAN exported the full ODE models in the Systems Biology Markup Language (Hucka et al., 2003) to ensure reproducibility, consistency, and interoperability.

### Parameter optimisation

We specified the parameter optimisation problem per dataset (drought or heat) in PETAB (Schmiester et al., 2021) and optimised the parameters of the ODE model to match experimental measurements on both the control and treatment condition using PyPESTO (Schälte et al., 2023) and AMICI (Fröhlich et al., 2021) for ODE integration. We started the ODE model with *t* = 0 being the measured initial module expressions and predicted the full time course data from there. Both conditions shared all parameters except for *u*(*t*) and *c*_*m*_(*t*). We minimised the negative log-likelihood objective function using the L-BFGS optimisation algorithm implemented in SciPy (Virtanen et al., 2020) and AMICI’s adjoint sensitivity method. For fitting, PyPESTO required the noise level for each module to calculate the negative log-likelihood. We set this value to 0.1, based on the 95% confidence interval estimated through bootstrapping of the mean per module. Parameter limits were chosen empirically to ensure the parameters of a best fit never hit their upper or lower bounds (Supp. Tab. S2), and all non-negative parameters were estimated on a log_10_ scale to aid exploring different orders of magnitude (Schälte et al., 2023). To maximally explore the optimisation landscape, models with random initial parameters were initialised 5,000 times in parallel, and waterfall plots were created to inspect if a global minimum was found (Supp. Fig. S9). In the heat dataset, the last two data points (*t≈* 10 and *t ≈*20) were oversampled fourfold to mitigate the effects of uneven sampling and ensure a more balanced representation across time points. Finally, to predict TF knockout effects *in silico* we set the parameter *β* of the associated edge(s) to 10^*−*6^. ODE fits and resulting parameters were stored using MLFLOW (Chen et al., 2020). Optimisation took approximately four days per dataset on a server equipped with two Intel(R) Xeon(R) E5-2640 v3 CPUs.

## Code availability

All source code isavailable at https://github.com/CropXR/BADDADAN.

## Author Contributions

BN, ADJvD and DdR conceived the initial idea. BN carried out the research and wrote the initial manuscript. MR, ADJvD, and DdR provided critical feedback. All authors have read and approved the final manuscript.

## Acknowledgments

The authors thank Tijmen den Butselaar for helping find the public data that was used to test BADDADAN, Elena Del Pup and Francesca Giaume for helpful discussions regarding the visualisation and biological interpretation of the results, and Jordi Alonso Esteve for input on the methodology.

## Supplementary Information

**Figure S1.**
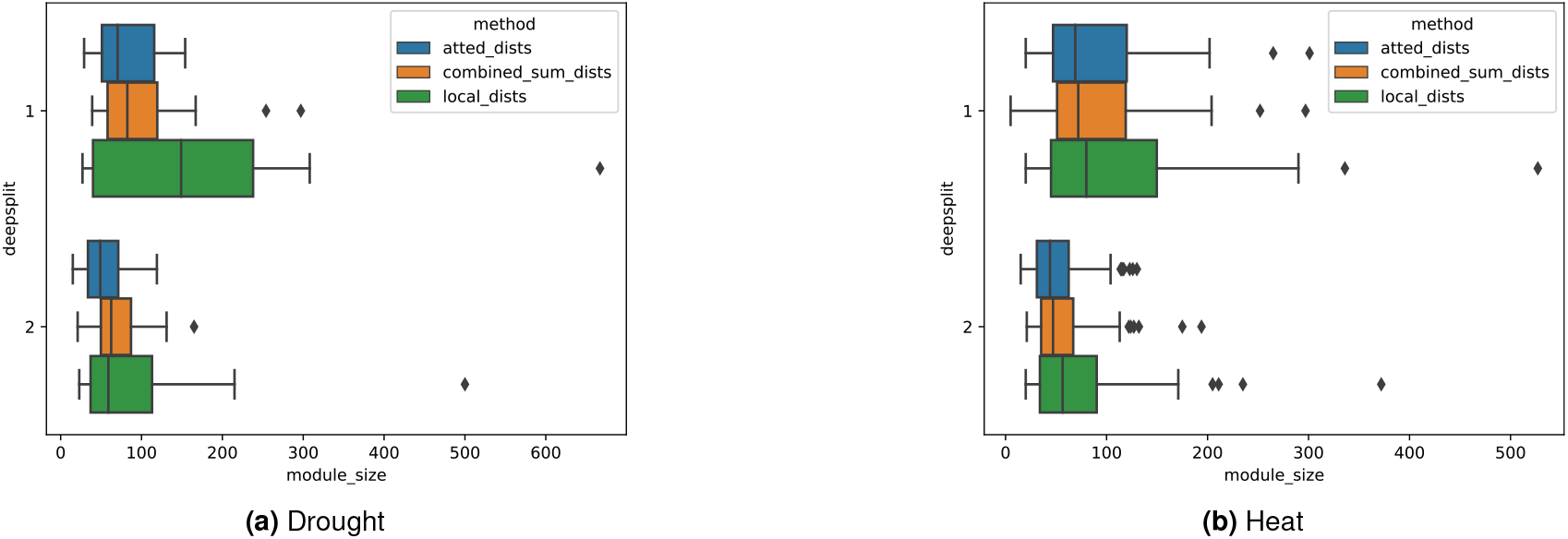
Module size distributions for both datasets at various values of the deepsplit parameter

**GO enrichment (Supp. Methods)**

**Table S1.** Number of modules per clustering. DS: deepsplit parameter

|  | Global | Combined | Local |
| --- | --- | --- | --- |
| Heat (all DS=1) | 52 | 52 | 42 |
| Drought (DS=2 for local, others DS=1) | 29 | 24 | 25 |

**Figure S2.**
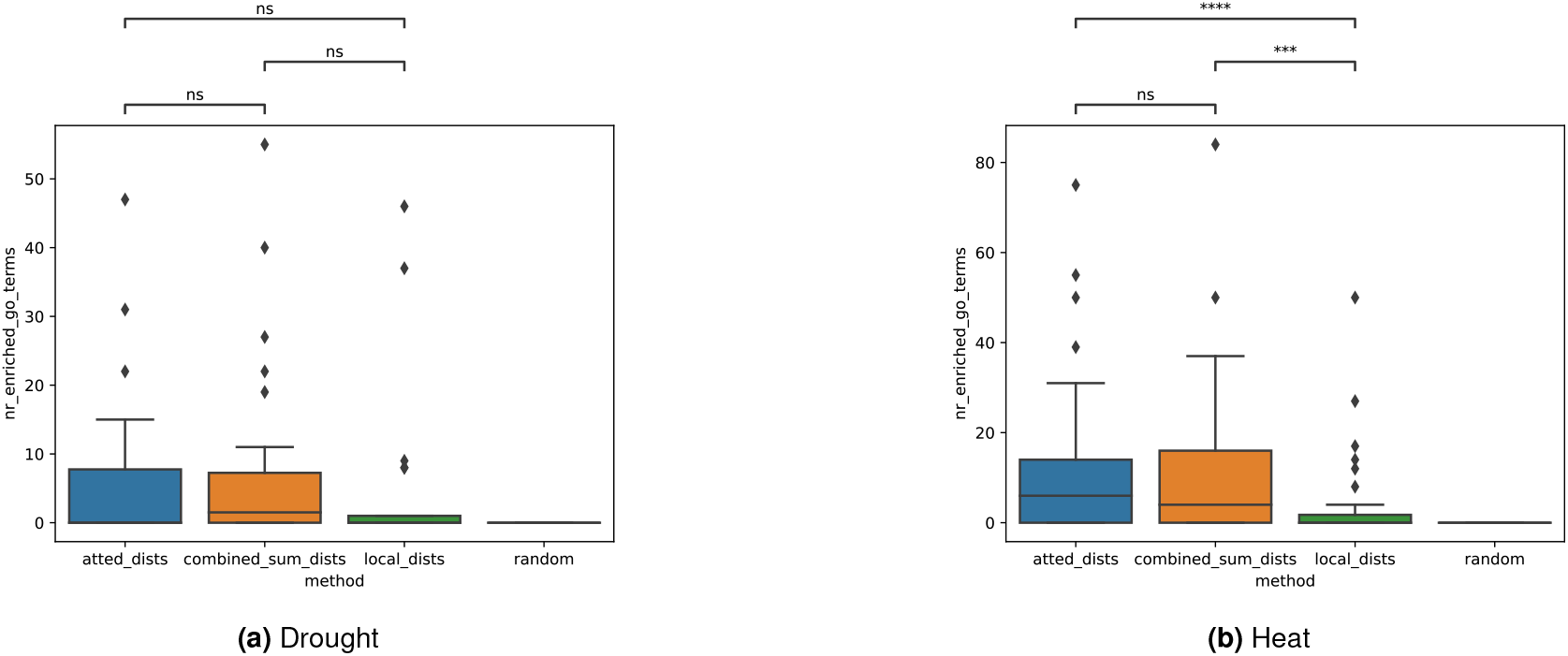
Number of enriched biological process GO terms per module. To ensure that the higher number of GO annotations did not result in unrelated—and thus hard to interpret—GO annotations, we also measured the semantic similarity of GO terms annotated per module. For this, we calculated the mean pairwise Wang (Wang et al., 2007) similarity score between all the enriched BP GO terms of a module using the Sswang function in GOAtools (Supp. Fig. S3).

**Figure S3.**
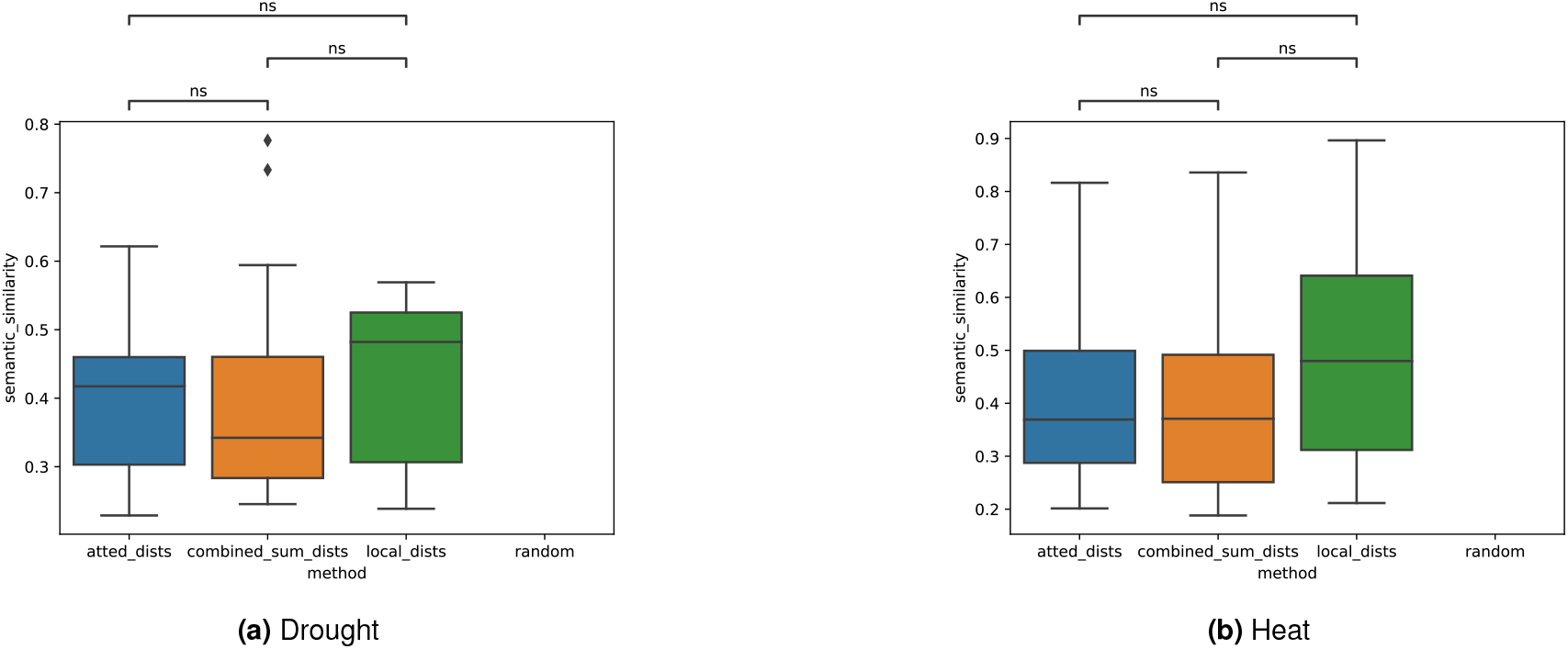
Mean pairwise semantic similarity between enriched BP GO terms per gene module

**Figure S4.**
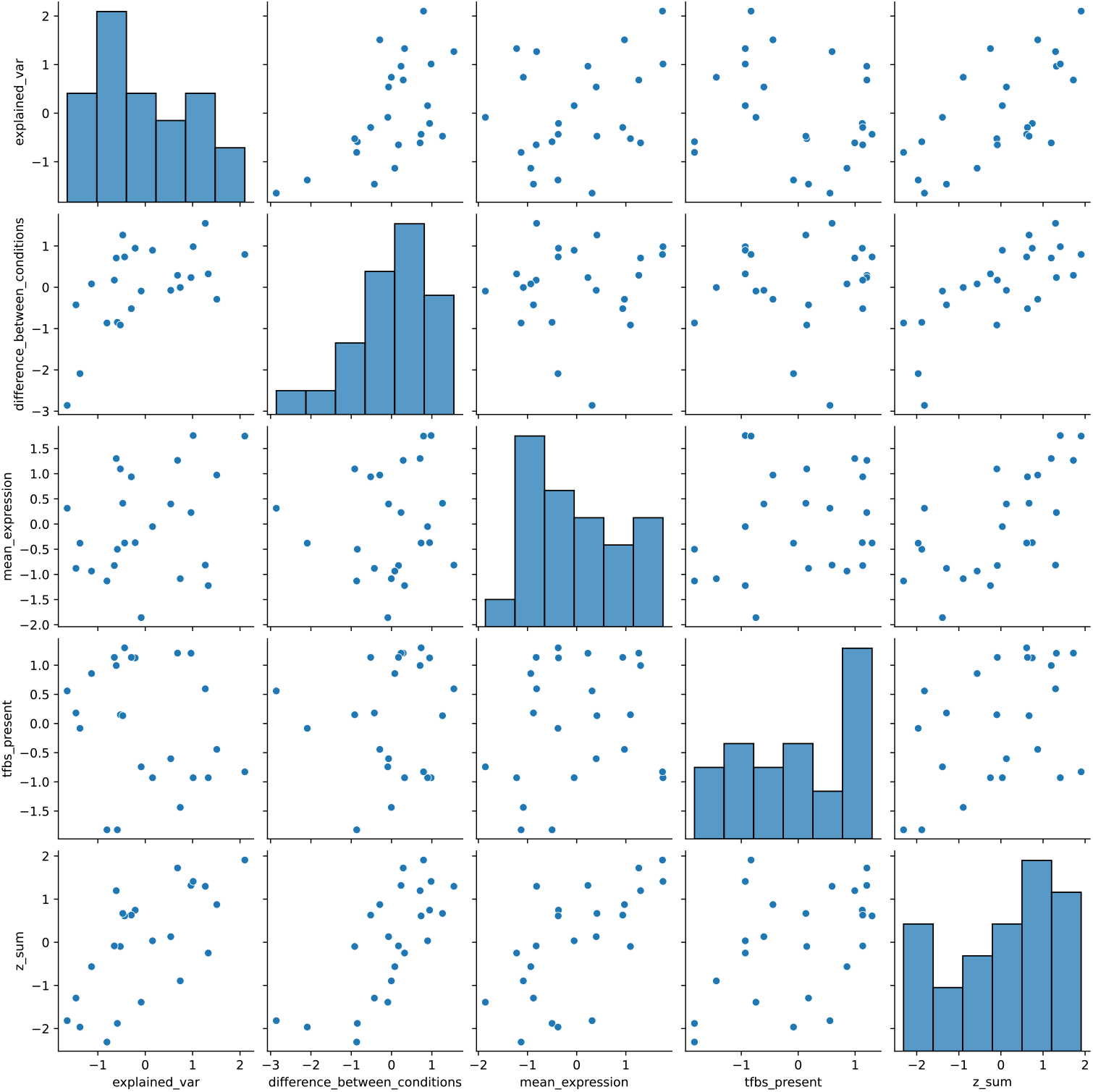
Z-scores of all modules in drought dataset

**Figure S5.**
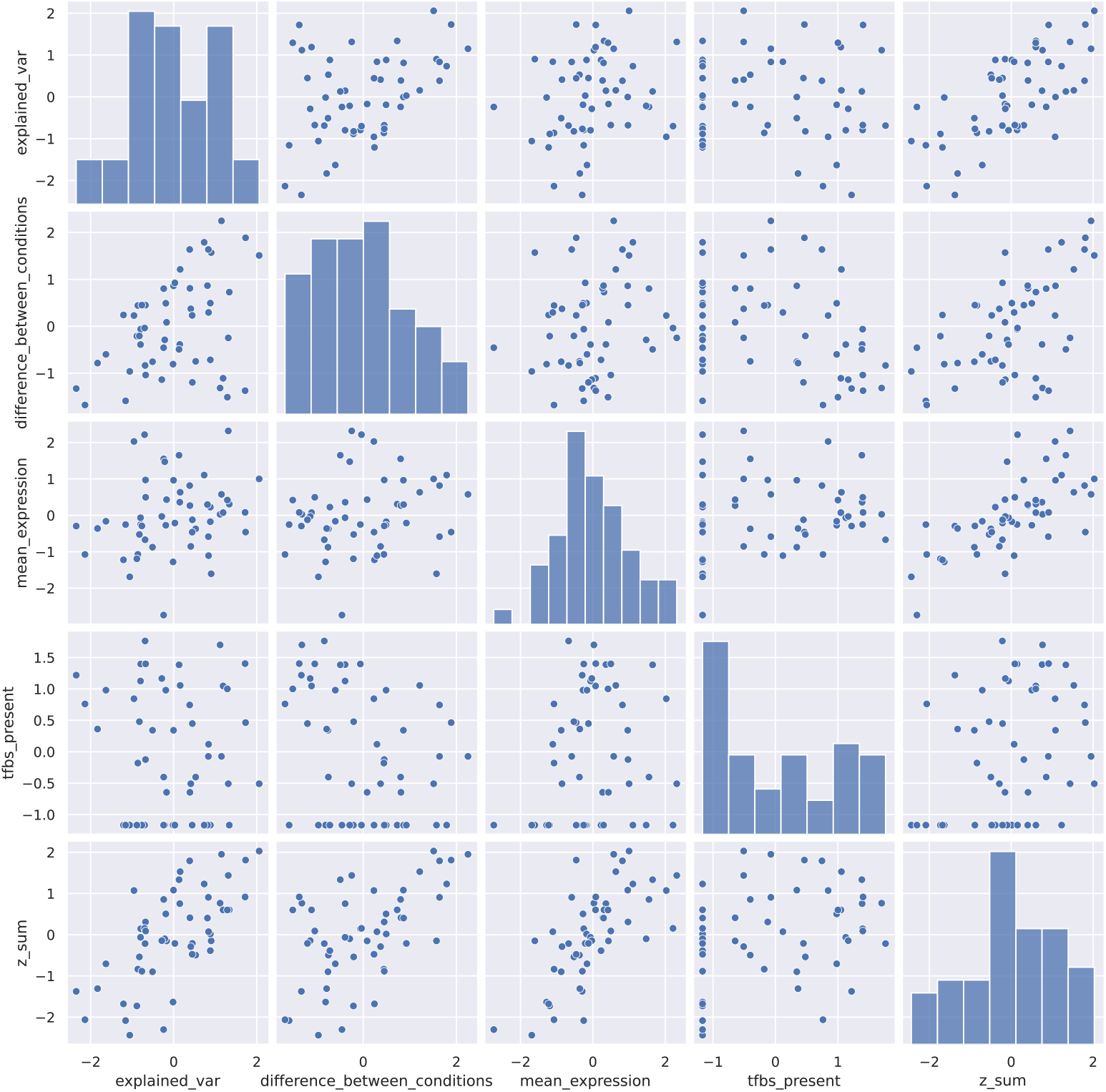
Z-scores of all modules in heat dataset

**Figure S6.**
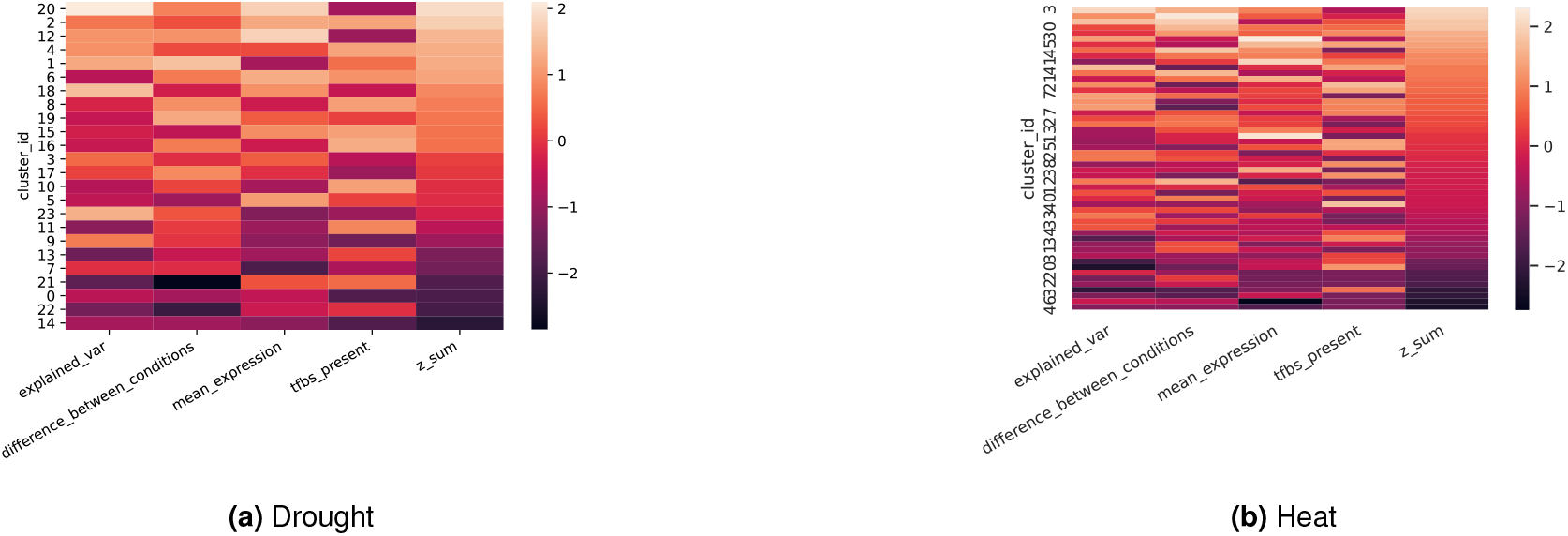
Breakdown of z-scores for all modules

**Figure S7.**
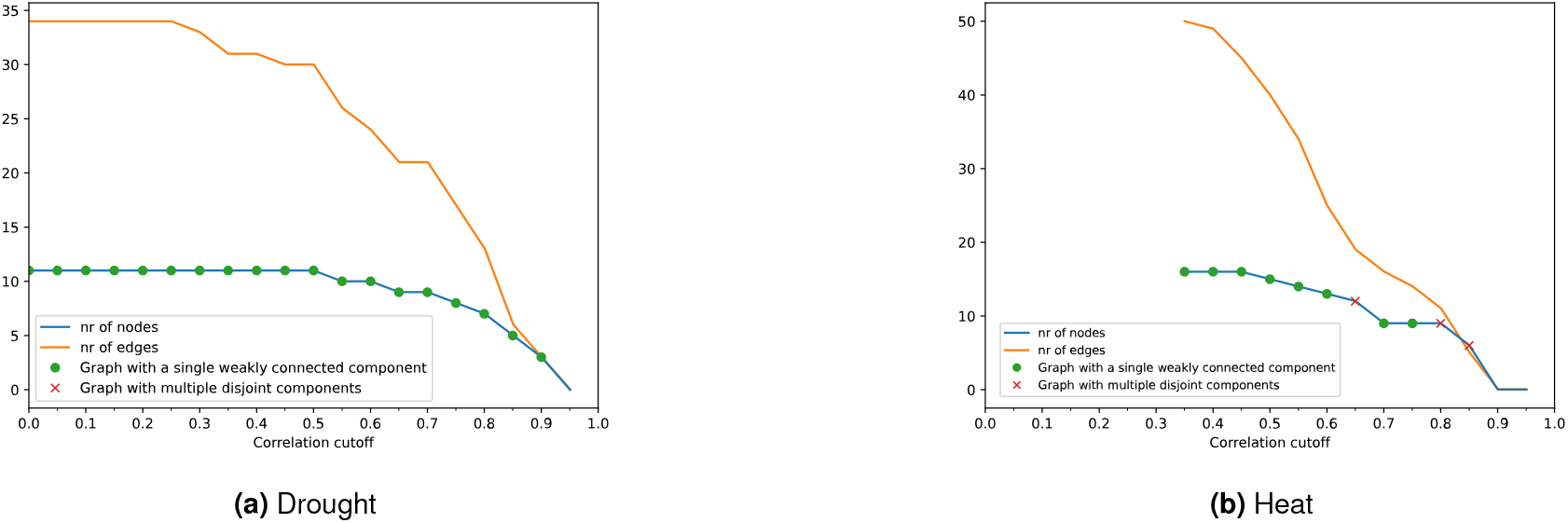
Characteristics of intermodular network for different correlation cutoffs to infer regulatory connections. When inconsistency between inhibition and activation between modules, no valid output was created because no graph could be inferred.

**Figure S8.**
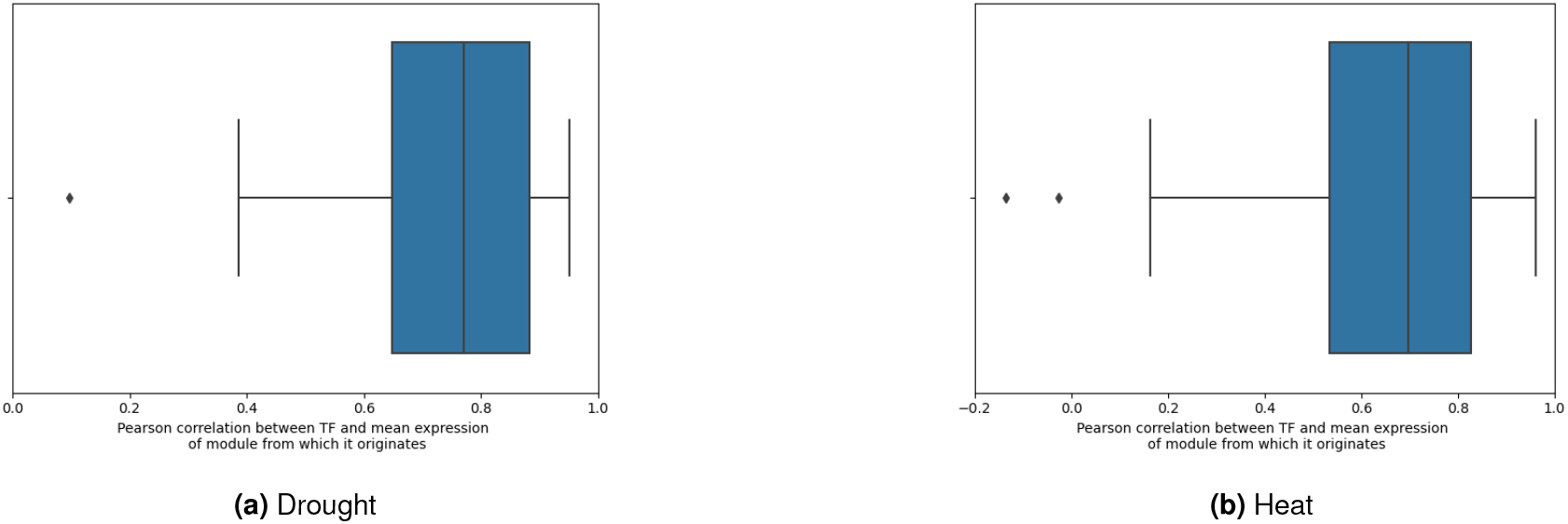
Pearson correlation between transcription factor expression levels and the mean expression of the module from which it is produced for both datasets.

**Table S2.** Parameter limits during fitting. *Parameter estimation scale* indicates if PyPESTO samples the parameters on a linear or log10 scale.

| Parameter | Parameter estimation scale | Minimum | Maximum |
| --- | --- | --- | --- |
| $\delta$ | lin | 0 | 50 |
| $\gamma$ | lin | -5 | 5 |
| $\beta$ | log10 | $10^{-6}$ | 1000.0 |
| $k$ | log10 | 0.1 | 10.0 |
| $a$ | log10 | $10^{-6}$ | 100.0 |
| $\phi$ | lin | $-\pi$ | $\pi$ |
| $b$ | lin | -10 | 10 |

**Figure S9.**
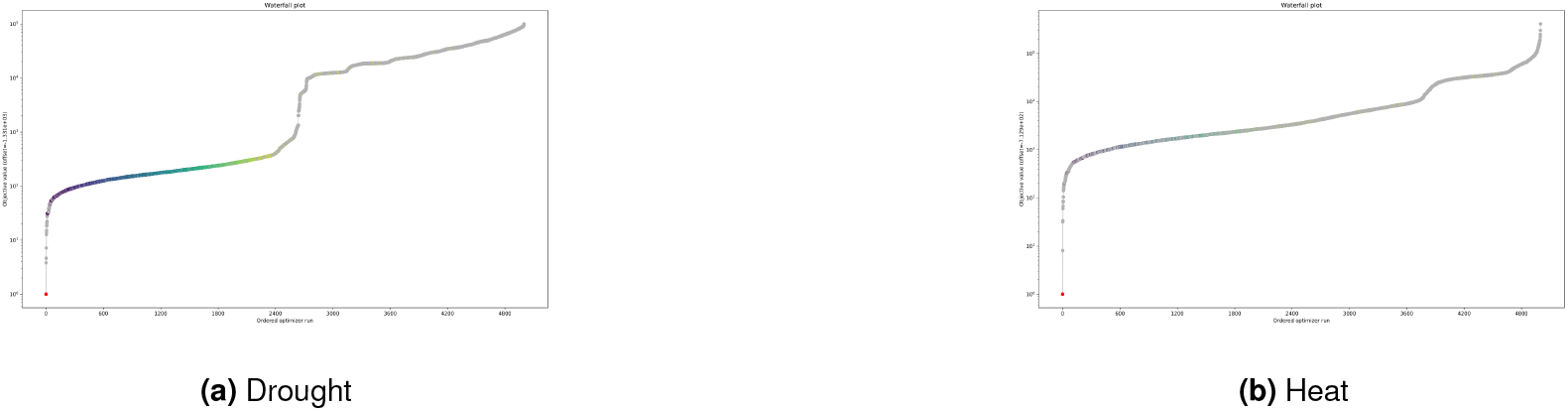
Waterfall plots created by PyPESTO. They show the objective values on the *y*-axis, with optimiser runs ranked from best to worst along the *x*-axis.

**Figure S10.**
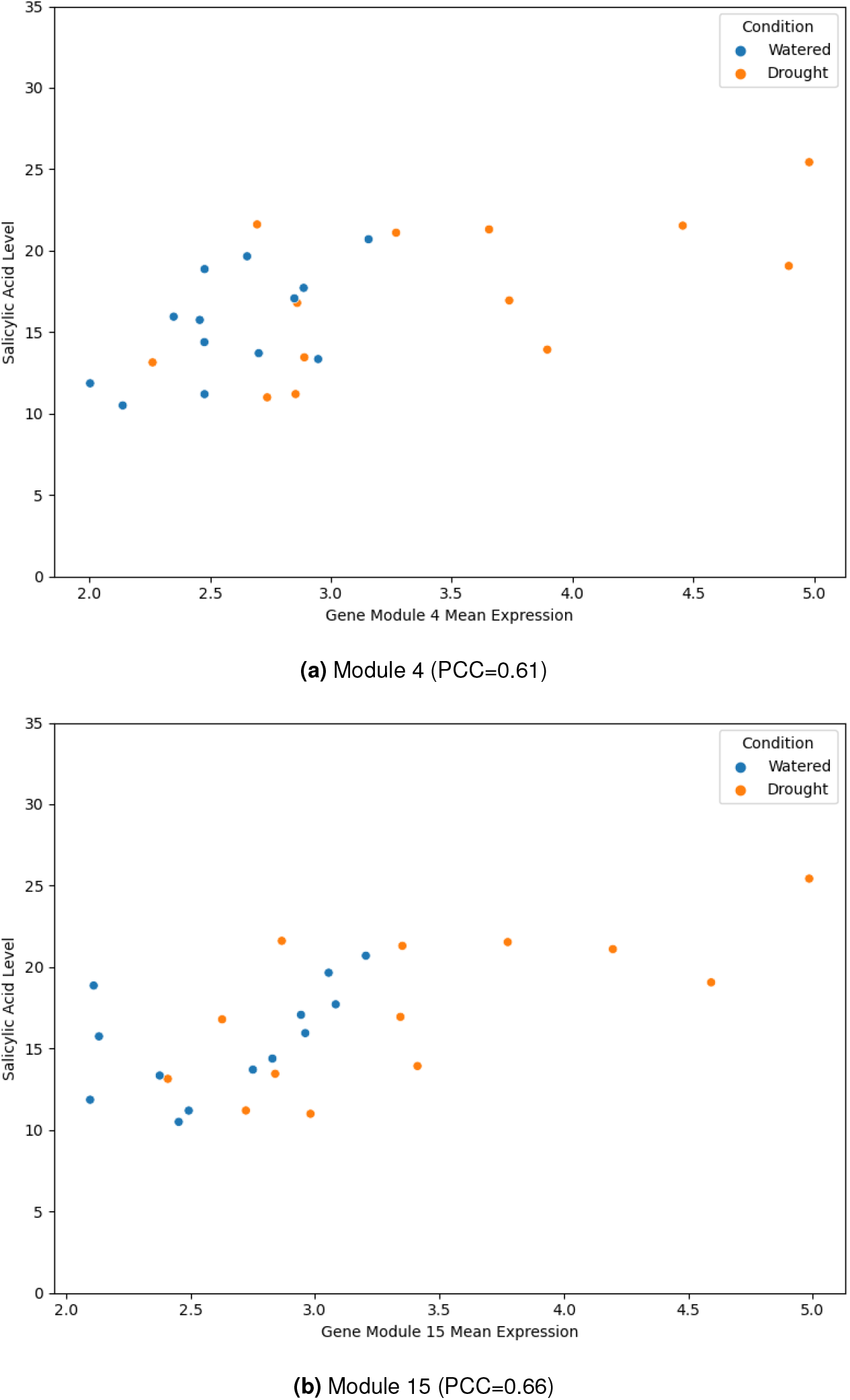
Pearson correlation of mean expression of gene module 4 and 15 (which were enriched for Salicylic Acid (SA)related GO terms) to abundance of SA in drought dataset. The SA levels were determined by LC-ESI-QToF MS metabolite profiling (Bechtold et al., 2016) and represent the mean peak area over 4 biological replicates.

**Figure S11.**
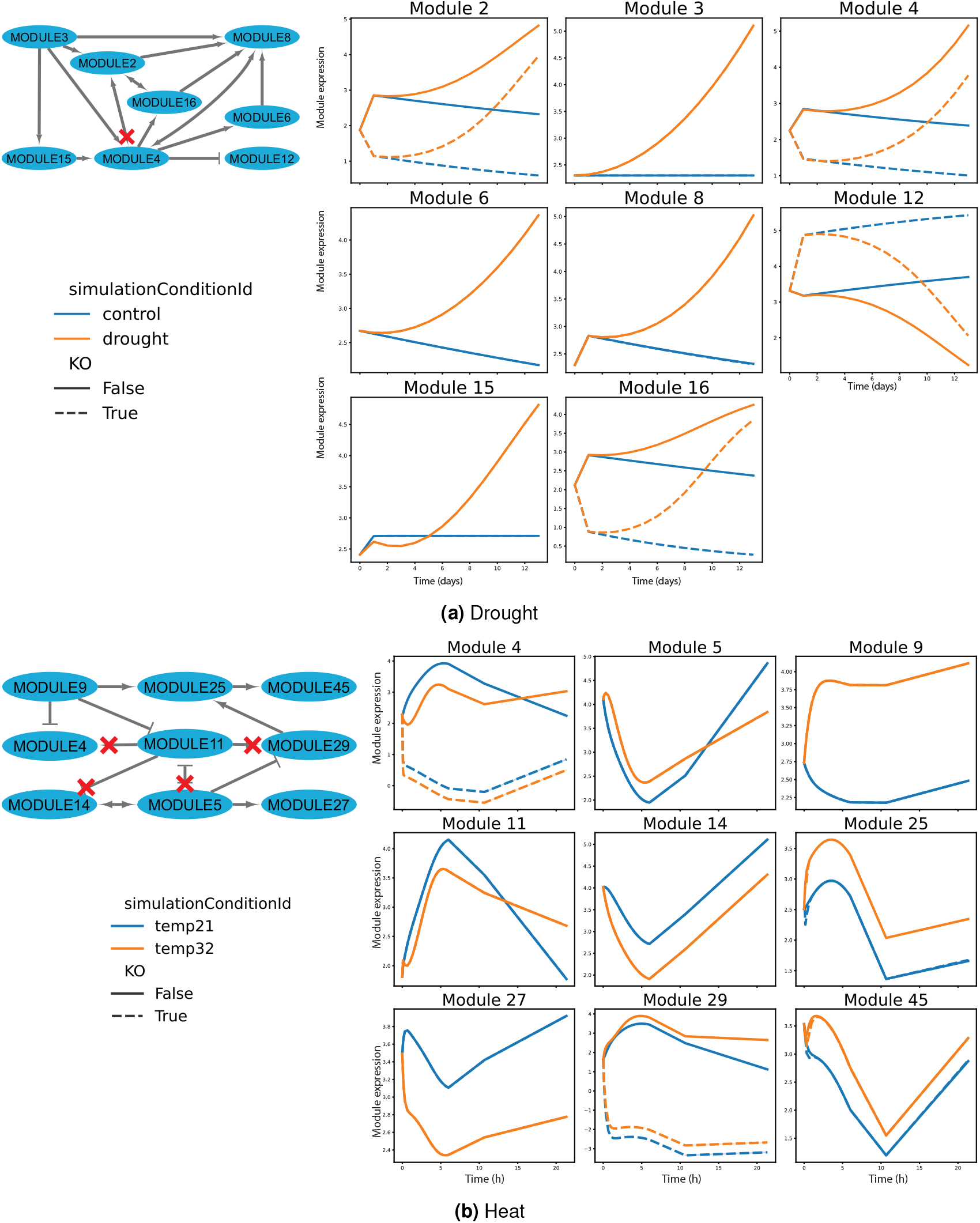
Knockout simulations of drought and heat. The red crosses indicate connections for which the knocked-out TF was responsible. **(A)** The knockout of CBF3 (AT4G25480) in drought. **(B)** The knockout of ABF1 (AT1G49720) in heat.

